# Topologically distinct and equivalent multiple efferent arterioles in the human glomeruli

**DOI:** 10.1101/2024.10.31.621302

**Authors:** Joshua Jing Xi Li, Jeremy Yuen Chun Teoh, Hei Ming Lai

## Abstract

We revisit the assumption that the human glomerulus consists of a capillary network with one afferent arteriole and only one efferent arteriole. Using three-dimensional tissue imaging, we unambiguously demonstrate the existence of multiple efferent arterioles via total reconstruction of a full human glomerular capillary network. Through graph network analysis, we found these multiple efferent arterioles can be topologically distinct or equivalent with reference to the global glomerular capillary network, and were strategically positioned with respect to the afferent arteriole. We also found the extraglomerular connections between two efferent arterioles may have significant contributions to the redistribution of intraglomerular blood flow. These new findings can provide new paradigms in the regulation and dysfunction of glomerular ultrafiltration, the crosstalk between intra- and extraglomerular factors, and the cross-coupling between nephrons.

## Main Text

The renal glomerulus is a contained microstructural unit with the specialised function of hydrodynamically driven ultrafiltration. The traditional textbook assumption was that glomeruli have a single afferent arteriole and a single efferent arteriole with an intervening capillary network. Such a description is based on the statistical sampling of many 2D images, which can be inaccurate since whole human glomeruli capillary network reconstruction in 3D has not been performed. Indeed, using the intravascular resin casting technique, Murakami et al revealed that multiple efferent arterioles might exist in the human renal glomerulus nearly 40 years ago ^1^. However, the study suffered from resin leakage, which may lead to artefactual connections to adjacent vessels or tubules that jeopardizes the validity of the finding, given the rarity of glomeruli with multiple efferents. In addition, whether these efferent arterioles are functionally distinct and hence responsible for their portion of blood flow from the glomerular capillary network is unclear. That is, the resin casting technique lacks the global topological information for distinguishing the connectivities of different efferent exit sites, and hence whether such multiple efferents carry hemodynamic significance.

The presence of multiple efferent arterioles within a subset of glomeruli can substantially change our understanding of renal function and ultrafiltration regulation. We decided to take a different approach to prove or disprove the presence of multiple efferent arterioles in the human glomerulus. Although many studies have attempted whole mouse kidney imaging for enumerating statistical changes under pathological and therapeutic conditions^2–6^, high-resolution 3D microstructural and capillary network analyses have not been attempted for individual glomeruli, especially for human samples. This is in part due to technical challenges in obtaining a complete map of the glomerular capillary network, namely, (1) the insufficient penetration of antibodies in the locally antigen-dense region, which leads to incomplete visualization of structures for analysis, (2) as a locally anchored structure, aggressive permeabilisation measures will likely distort or even crack the delicate structures within the glomerulus. Finally, (3) the need for fortuitously sectioned renal glomeruli that fall within the working distance of an oil objective under a confocal microscope, which is typically restricted to ∼150μm, prohibiting the imaging of large human glomeruli. Missing any blood vessels, due to cutting artefacts or limits in imaging depths, would introduce uncertainties in the connectivity of the capillary network. Such reconstruction is essentially all-or-none. To date, there has been no fully reconstructed capillary network for the human renal glomerulus. In this study, we thus set our goal to provide the first fully reconstructed human glomerular capillary network graph and demonstrate the topological distinctiveness of the multiple efferents both inside and outside of the glomerular globe.

After screening around 50 glomeruli from 4 individuals (2 males and 2 females), we found 4 glomeruli (**Supp Fig 2, 3**) from two individuals to have multiple efferents, and only one of them can be fully imaged for global network analyses. These 4 glomeruli multiple efferents were found in the 2 male subjects (3 glomeruli from the 1st subject, aged 79, and 1 glomerulus from the 2nd subject, age 58). After manual tracing and segmentation, the capillary network can be represented as a graph *G =* (*V, E*), where capillary branch points are nodes {*v*_*1*_, *v*_*2*_, … *v*_*n*_ ∈*V*} and the capillary segment between the branch points are edges {{*v*_*i*_, *v*_*j*_} | *i* ≠ *j*} ∈*E* of the graph (**Fig. 1** and **Supp Video 1**, please see also **Methods**). The weights of the edges are given as the lengths of the capillary segment, or as their inverse which is proportional to the conductance of fluid flow by the Hagan-Poiseullie equation. The full network consisted of 454 branch points and a total capillary length of 20,235.20 μm, of which 18,010.96 μm (89.01%) are intraglomerular. The full image stack revealed a single afferent arteriole entry site (denoted as Aff_1_) identified based on its characteristic muscular wall and 4 efferent vessels exit sites from the glomerular globe, which we denote as Eff_1_, Eff_2a_, Eff_2b_, and Eff_3_. Interestingly, Eff_2a_ and Eff_2b_ are promptly re-connected by the traced extraglomerular vessels, while the extraglomerular vessel trajectories of Eff_1_, Eff_2_ and Eff_3_ diverge. All four exit sites are spatially located close to the juxtaglomerular region, suggesting that classical regulatory mechanisms mediated by the juxtaglomerular apparatus apply to them.

Anatomically, Eff_1_ has the widest opening and is closest to the juxtaglomerular apparatus, while Eff_3_ exit the glomerulus at very close proximity to Aff_1_ and appears to have a circular muscle lining. How are the distinct efferent vessels positioned within the glomerular capillary network? We partitioned the glomerular capillary network into two lobules with the sparsest cut, which is based on the graph’s normalised conductance-weighted Laplacian matrix. We also assigned each point of the network to its nearest efferent exit site, which we refer to as the efferent exit site receptive domains, revealing that each of the 4 efferent exit sites receives blood flow from a substantial proportion of the intraglomerular capillary network, suggesting that they provide non-redundant exit sites for the intraglomerular blood flow (**Fig 1a**). Using this classification scheme, we note that Aff_1_ is located at the interface between the two lobules - a strategic position within the glomerular network. The exit sites are also clearly divided in their lobule subnetwork assignment, with Eff_2a_ and Eff_2b_ deeply located in lobule 2, and Eff_1_ and Eff_3_ deeply located in lobule 1. By comparing lobules and domains, we found lobule 2 is constituted by mostly Eff_2a/b_ domains and a small portion of the Eff_1_ domain. Conversely, Eff_2_ domains are exclusively located in lobule 2 (**Fig. 1b**).

**Figure 1.**
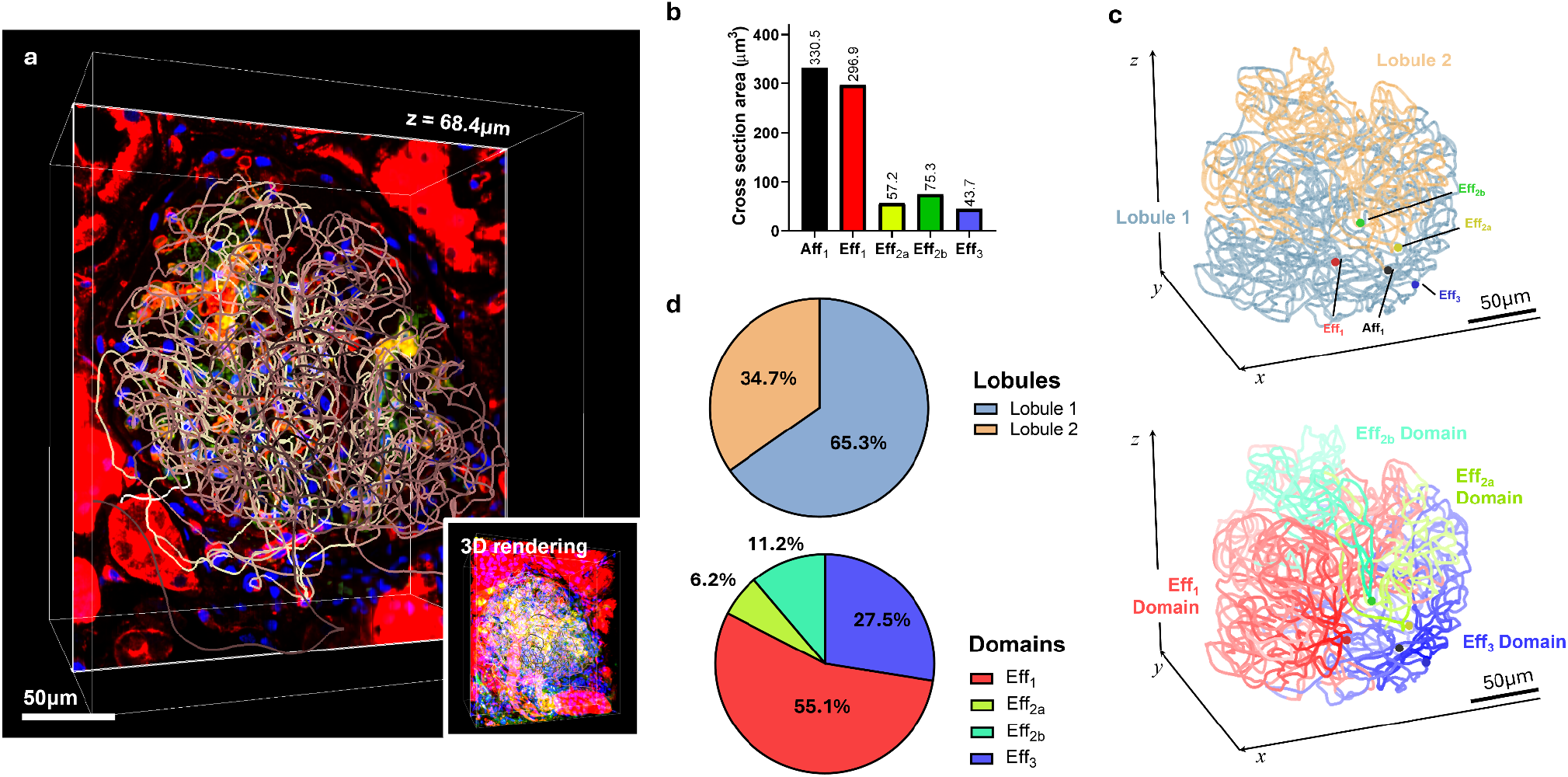
Organization of a human glomerular capillary network with multiple efferents vessels. **a**. 3D image volume of the human glomerulus with multiple efferent vessels. Only a z-slice is shown in the larger image for clarity, the inset shows the fully rendered image volume. The full tracing has been overlaid in the same 3D space. **b**. Cross-section areas for the afferent arterioles and the 4 efferent exit sites. **c**. Complete views of the intraglomerular capillary network with colouring according to lobules and domains. The Aff_1_ and Eff exit sites have been highlighted as a dot and coloured accordingly. **d**. The proportion of capillary segments (in length percentages) for each lobule and domain.

How are these four exit sites interrelated via the glomerular capillary network? To address this question, we examined all the pairwise shortest paths between the afferent and efferent vessels (**Fig. 2a, Supp Fig 4**). We found that the shortest paths connecting Eff_2a/b_ and Eff_3_ always pass through Eff_1_ or Aff_1_. Specifically, we found the shortest path *P*_**min**_(Eff_2a_ ↔ Eff_3_) = *P*_**min**_(Eff_2a_ ↔ Aff_1_) ∪ *P*_**min**_(Aff_1_ ↔ Eff_3_), and *P*_**min**_(Eff_2b_ ↔ Eff_3_) = *P*_**min**_(Eff_2b_ ↔ Eff_1_) ∪*P*_**min**_(Eff_2b_ ↔ Eff_1_) (**Fig. 2a, b**). Hence, Eff_2_ and Eff_3_ are topologically separated by the glomerular capillary domains of Eff_1_ and Aff_1_. These results indicate that topologically distinct and equivalent exit sites exist for a human glomerulus. In addition, we also note that *P*_**min**_(Aff_1_ ↔ Eff_3_) and *P*_**min**_(Aff_1_ ↔ Eff_2a_) are distinctively shorter than others. Taken together, the arrangement of exit/entry sites of this glomerulus forms a symmetric pattern (**Fig. 2c**). Since hydrodynamic pressure drop is inversely proportional to the length of a vessel, such observation suggests that capillary segments topologically close to the Eff_3_ and Eff_2a_ exit sites are particularly sensitive to back pressure effects on their efferent vessels.

**Figure 2.**
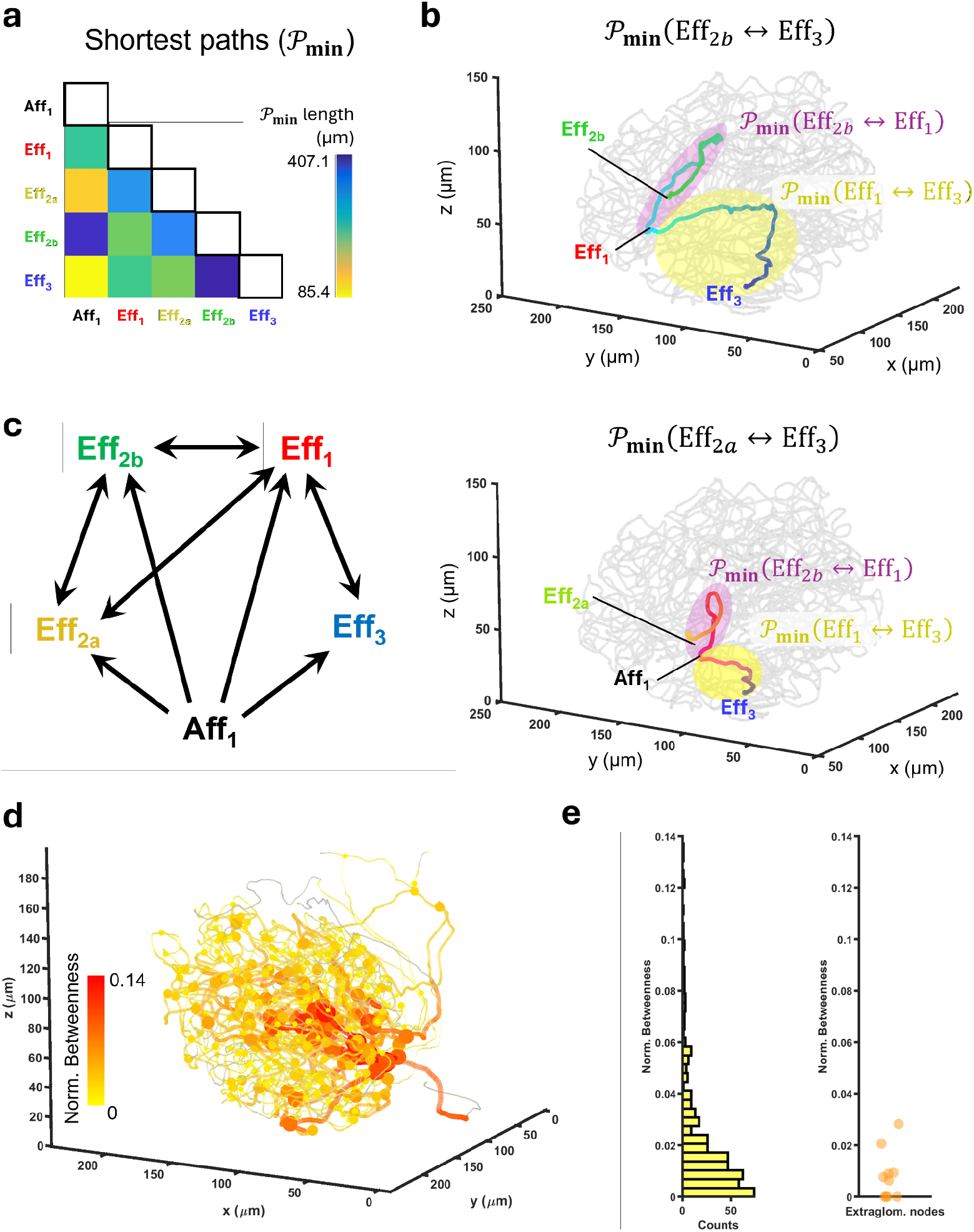
Positioning of the multiple efferents in the global capillary network. **a**. Computed pairwise shortest paths between Aff_1_ and each efferent vessel exit site. **b**. Visualization of the shortest paths that are the complementary concatenation of other shortest paths between the exit/entry sites. **c**. Abstracted connectivity map between the exit/entry sites based on shortest path analysis. **d**. Plot of nodes with their size and colour according to their betweenness centrality. **e**. The left histogram shows the normalized betweenness centrality of all 454 nodes, while the right plot shows only the extraglomerular nodes’ normalized betweenness centrality data.

We also found there exist direct, multiple extraglomerular connections between Eff_2a_ and Eff_2b_. Computing the weighted betweenness centrality of each capillary segment revealed that such extraglomerular connections have strong total network connectivity contributions (**Fig. 2d**), even higher than many intraglomerular capillary segments (**Fig. 2e**). This result suggests that a significant proportion of blood flow can be directed from one glomerular capillary locus to another via these extraglomerular connections. Equivalently, if there is an appropriate pressure difference within the intraglomerular capillary network, such topological arrangement suggests a high probability that blood can flow out of Eff_2a_ and re-enter via the “exit” site Eff_2b_, and vice versa.

## Discussion

The single nephron glomerular filtration rate (SNGFR) is dependent on the ultrafiltration coefficient (*K*_*f*_), the transluminal hydraulic (*ΔP*) and oncotic pressures (*Δ*Π), and the glomerular blood flow (*Q*), given as SNGFR = *K*_*f*_ (*ΔP - Δ*Π)^10^. The SNGFR can also be expressed as the sum of all capillary segments’ ultrafiltration contributions, with their own sets of *K*_*f*_, *ΔP, Δ*Π and *Q* determining the single capillary segment filtration rate (SCSFR). At a given afferent hydraulic and oncotic pressure and blood flow, a maximal SNGFR will occur when the distribution of transluminal ultrafiltration pressure and blood flow is matched to the filtration surface area fraction of each capillary segment - i.e., an optimal perfusion-filtration matching. The discovery of multiple, topologically distinct efferent arterioles allows a much more nuanced local regulation of SCSFR, where a higher back pressure of an efferent arteriole will cause higher transluminal hydraulic pressure but less blood flow for capillary segments proximal to it. Solving the dynamical regulation of SCSFR may be key to understanding the pathological changes and progression underlying segmental glomerulonephritis conditions.

Through network analysis, we have revealed the organization of multiple efferent arterioles in the human glomerulus. Specifically, the Laplacian matrix-based sparsest cut provides information on the network’s global topology, which aims to cut the network into two components by cutting the minimal number of edges with preferably the weakest conductances. Meanwhile, domain assignment based on the nearest efferent exit site is based on structural assessment, based on how the blood flow is likely distributed across the different exit sites. The combined analyses based on lobule partition, domain assignment and shortest path analyses suggest the segregation of Eff_2a/b_ from both Eff_1_ and Eff_3_ is structurally and topologically consistent. Concerning the abstracted organization of this glomerulus in **Supp Fig. 5**, there seems to be a hierarchy of lobules with potentially distinct functions. Eff_1_ and its domain perhaps contribute to the majority of the ultrafiltration and receive the most blood flow by this glomerulus, while the domains of Eff_2_ and Eff_3_ are “accessory” pathways that may serve some regulatory or sensing functions. Their topological segregation suggests they may operate independently. For example, back pressure on Eff_3_ due to intra- or extraglomerular factors would be expected to affect the Eff_1_ domain first, before propagating further to Eff_2_ efferent lobules, and vice versa.

The presence of multiple efferent arterioles also opens up an intriguing possibility where multiple nephron functions can be coupled with glomeruli acting as the hub of regulation. If the different efferent arterioles of the glomerulus drain into different vasa recta, the same glomerulus could sense and integrate the back pressures built up across multiple nephrons. Humoral factors released by a single juxtaglomerular apparatus can also be broadcasted to different nephrons. Hence, multiple efferents in a single glomerulus can provide the structural basis for cross-nephron coupling in signalling and hemodynamic functions.

Finally, the significance of extraglomerular connections between the efferent exit sites has also been established through centrality analysis, which may be an important transit pathway for intraglomerular capillary blood flow. Apart from the possibilities of serving as a non-filtrating blood flow shunt, pressure-transducing functions, or cellular migration conduits, it may also hint at unknown mechanisms on how intra- and extraglomerular cellular and molecular mechanisms can cross-influence each other in healthy and pathological situations. This can be an intra- and extraglomerular crosstalk channel in addition to the gap junctions between intraglomerular and extraglomerular mesangial cells^11^.

We acknowledge that this study is limited by only finding 3 glomeruli with multiple efferent arterioles across two individuals, which warrants further systematic studies on whether this is a ubiquitous phenomenon. There are also many open questions, such as the distribution of these multi-efferent human glomeruli, their vascular connectivity, and the revised structural-functional relationships in light of such discovery. Their presence and implications in pathological conditions remain to be addressed. In summary, 3D histology with complete mapping of the glomerular capillary network has unambiguously confirmed that multiple efferent arterioles can exist in the human renal glomerulus. Our study highlights the need for systematic, assumption-free 3D histological investigations of structures considered to be well-known, which can lead to an unexpected structural variable that can substantially change our physiological and pathological mechanistic formulations.

## Methods

Human tissues were obtained from healthy cadavers. Ethical approval was obtained from The University of Hong Kong / Hospital Authority Hong Kong West institutional review board with the waiver of the written consent requirement (reference number: UW 24-406). Tissues were obtained within 2 weeks of post-mortem delay and fixed in neutral buffered formalin for 1 week at room temperature, washed in 1x PBS and stored at 4°C until use. For surgical tissues, samples were obtained from total nephrectomy due to renal cell carcinoma treatment. We dissected out the grossly normal tissue volumes far from the tumour region, which were then promptly fixed in neutral buffered formalin for 1 week at room temperature, washed in 1x PBS and stored at 4°C until use.

The autopsy kidneys were dissected and perfused with a CM-DiI (Invitrogen, C7000) solution to obtain a complete vascular configuration of the glomerulus. The perfusion solution was prepared as previously described^7^. The perfused kidney cortex was then dissected and cut into ∼10mm x 5mm x 1mm-thick sections by hand, post-fixed in 4% paraformaldehyde in 1x PBS at room temperature (r.t.) overnight, washed two times with 1x PBS at r.t. twice for 30 minutes, and proceeded to tissue delipidation using dichloromethane/methanol mix and the INSIHGT protocol for 3D immunostaining^8^. INSIHGT was performed with 2 days of macromolecular probe labelling using rabbit anti-CD31 (Abclonal, A0378, at 10μg/ml), AlexaFluor 647-labelled Donkey anti-rabbit Fab fragments (Jackson Immunoresearch, 711-607-003, at 10μg/ml), and Dylight 649-labelled Lycopersicon esculentum lectin (VectorLabs, DL-1178-1, at 10μg/ml) in 1ml of 1x INSIHGT solution A. 5μl pre-mixed DAPI/INSIHGT solution C was added for nuclear staining at 2.5μg/ml DAPI final concentration in INSIHGT solution A. For nephrectomy specimens, perfusion of lipophilic tracer was impossible and hence a mouse anti-aSMA primary antibodies (Progen, 690001, at 10μg/ml) plus AlexaFluor 594-labelled Donkey anti-mouse Fab fragments (Jackson Immunoresearch, 715-587-003, at 10μg/ml) where added for visualizing the smooth muscle cells around the arterioles. The stained tissue was cleared using the benzyl alcohol/benzyl benzoate (BABB) method^9^. Imaging was performed using a Leica SP8 confocal microscope with a 40x oil objective (Leica HC PL APO 40X/1.30 Oil CS2) with excitation at 405nm, 561nm and 649nm. We used ultra-thin cover glass-bottomed confocal dishes with #0 cover glass (0.085-0.115 mm, from CellVis, D35-20-0-N), which provides 50μm more working distance and increased the success rate of finding three dimensionally intact whole glomeruli. An overview of the tissue processing procedure has been summarized in **Supp Fig 1a**.

The image stacks were first aligned to correct for any translational drifts, then segmented by hand in ImageJ, where a dot is placed near the centroid of the vessel lumen in each image slice. These dots connected throughout the image stack will become the tracing of the capillary network of the glomerulus. The tracing was then cleaned, skeletonised, converted to a graph (**Supp Fig 1b**), and analysed using custom-written scripts in MATLAB (Mathworks 2024).

We denote the glomerular capillary graph as a weighted graph *G =* (*V, E*), where capillary branch points are nodes {*v*_*1*_, *v*_*2*_, … *v*_*n*_ ∈*V*} and the capillary segment between the branch points are edges {{*v*_*i*_, *v*_*j*_} | *i* ≠ *j*} ∈*E* of the graph. To preserve the tortuous trajectories of the capillaries, multiple vertices *v*_deg=2_ = {deg(*v*) = 2} ⊆ *V* were preserved for the skeleton-to-graph conversion process, with a tabulated function *f* : *V* ↦ ℝ^*3*^ that maps their position back to the physical space. When computing for the shortest paths, the edges’ weights between each node take values as their Euclidean lengths 𝓁_*e*_, while for topological analysis, the edge weights take values as length-dependent conductance due to *c*_*e*_ ∝ 1*/*𝓁_*e*_. The normalized weighted Laplacian of *G* is defined as *L = I - D*^*-*½^*AD*^*-*½^, where *I* is the identity matrix of size *n × n, D* is the degree matrix, and *A* is the *c*_*e*_-weighted adjacency matrix of the graph. For the set of eigenvalues of *L* as λ_1_ ≤ λ_2_ ≤ … ≤ λ_*n*_ that are solutions of the equation *L***x** = λ**x**, with the vector **x**’s entries as *x*_*n*_, we define lobule 1 vertices as {*v*_*i*_ | *x*_*i*_ ≥ 0} ∈*V*_lobule 1_ and {*v*_*j*_ | *x*_*j*_ < 0} ∈*V*_lobule 2_ with (*V*_lobule 1_ ∪*V*_lobule 2_) = *V* for the case λ_2_ (i.e., the Fielder eigenvalue). We also assigned each vertex to its nearest efferent exit site, based on **arg min**_*s*∈*S*_[*d*_*G*_(*v, s*)] where *s* ∈*S* is the set of all efferent exit sites, and *d*_*G*_ is the distance based on the graph embedding of the glomerular capillary network. Betweenness centrality was computed as [2/(*n*-1)(*n*-2)]*×*∑_*s≠t≠v*_ *ρ*_*st*_(*v*)/*ρ*_*st*_, where *ρ*_*st*_(*v*) is the total number of shortest paths that pass through node *v*, and *ρ*_*st*_ is the total number of shortest paths between node *s* and *t*, and *n* is the total number of nodes in the graph. In this case of a conductance-weighted graph matrix, *ρ*_*st*_ is usually 1.

## Acknowledgement

We would like to express our deepest gratitude towards the tissue donors that enabled this study. We also thank Ms Carmen Chan and Dr Cathy Chan for logistics and administrative support.

## Data availability

The tracing data of the glomerulus shown in the figures have been deposited on CodeOcean (https://codeocean.com/capsule/2358628/tree). The source code that provides topological analysis and plotting of all the figures was provided in the same link.

