## Supplementary material for "Topologically distinct and equivalent multiple efferent arterioles in the human glomeruli": Supp Fig 1-5

**Supplementary Materials**

Supplementary Figures 1-5
Supplementary Video 1 caption

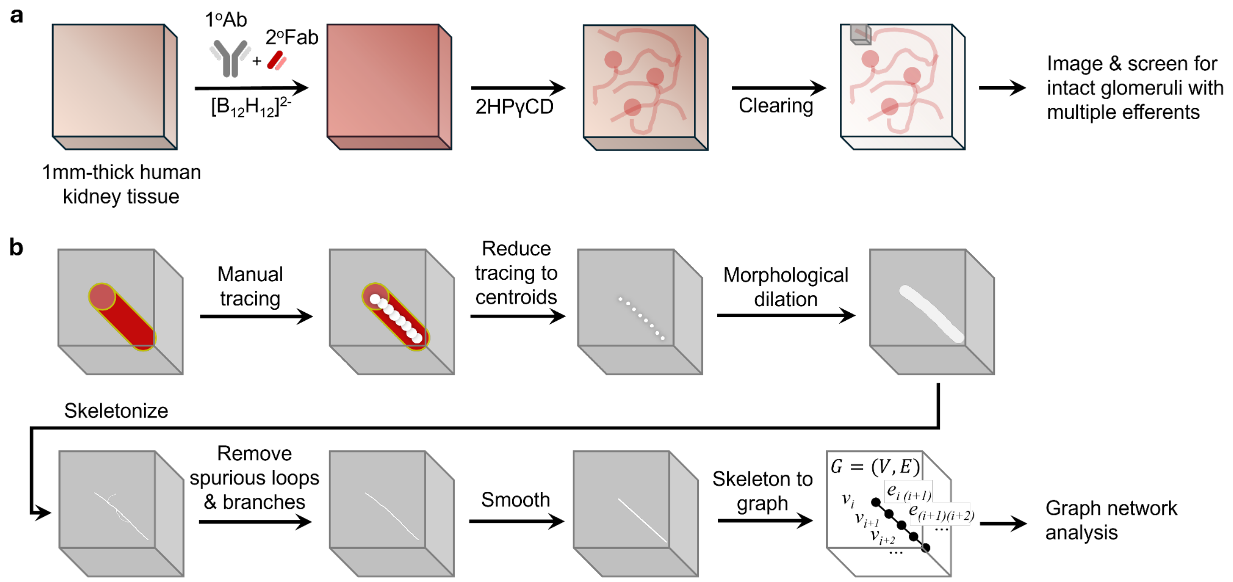


**Supp Fig 1. Overview of experimental method and image analysis for this study. a**. Tissue processing of 1mm-thick human kidney tissues for screening for glomeruli with multiple efferents. 1°Ab: primary antibody, 2°Fab: secondary antibody Fab fragment. **b**. Image analysis pipeline converting glomerular vasculature to a graph network via manual tracing and skeletonization.

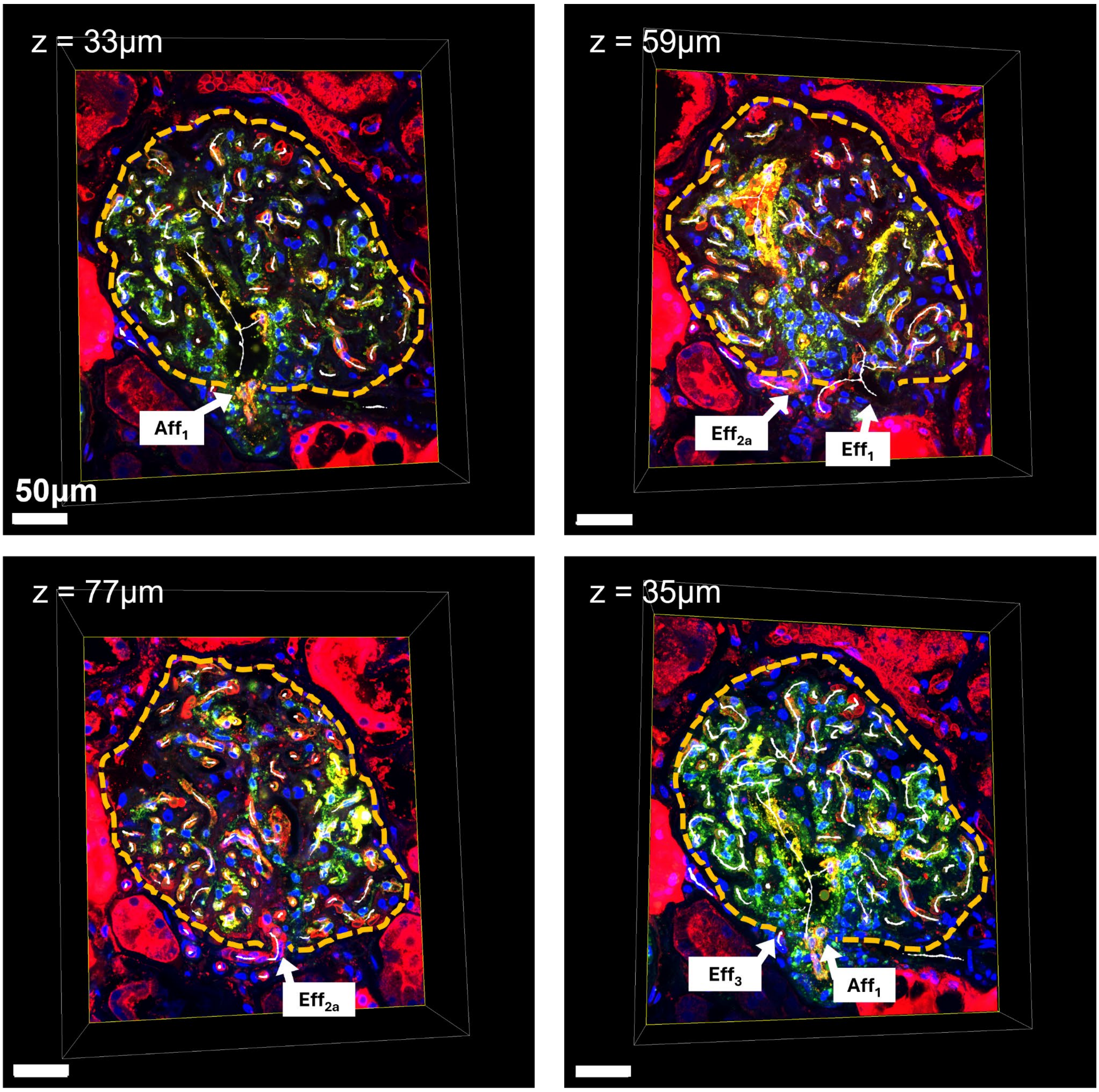


**Supp Fig 2. $Aff_1$ and the 4 Eff exit sites as shown in different z-slice levels of the 3D volume.** In red: CD31 and *Lycopersicon esculentum* lectin staining, in green: CM-DiI tracing, in blue: DAPI staining.

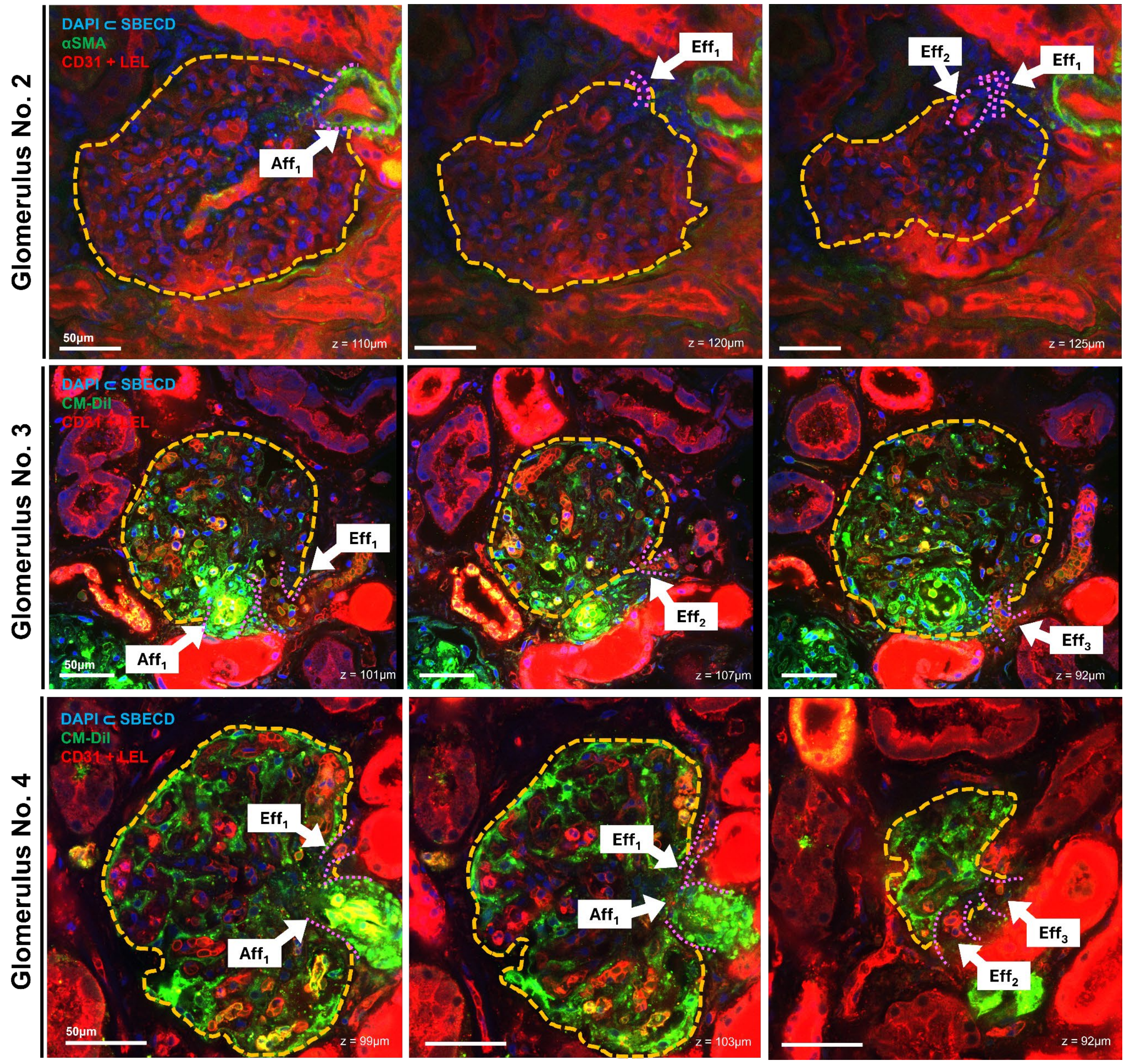


**Supp Fig 3.** Other human glomeruli with multiple efferent arterioles were found. The numbering of efferent exit sites (Eff) is arbitrary. Glomerulus No. 2 is from a total nephrectomy specimen devoid of renal cell carcinoma, while Glomerulus No. 3 and No. 4 are from an autopsy kidney specimen. αSMA: alpha-smooth muscle actin immunostaining CD31 + LEL: CD31 immunostaining and *Lycopersicon esculentum* lectin histochemical staining, CM-DiI: CM-DiI perfusion staining, DAPI⊂SBECD: nuclear staining with DAPI complexed by sulfobutylether-β-cyclodextrin.

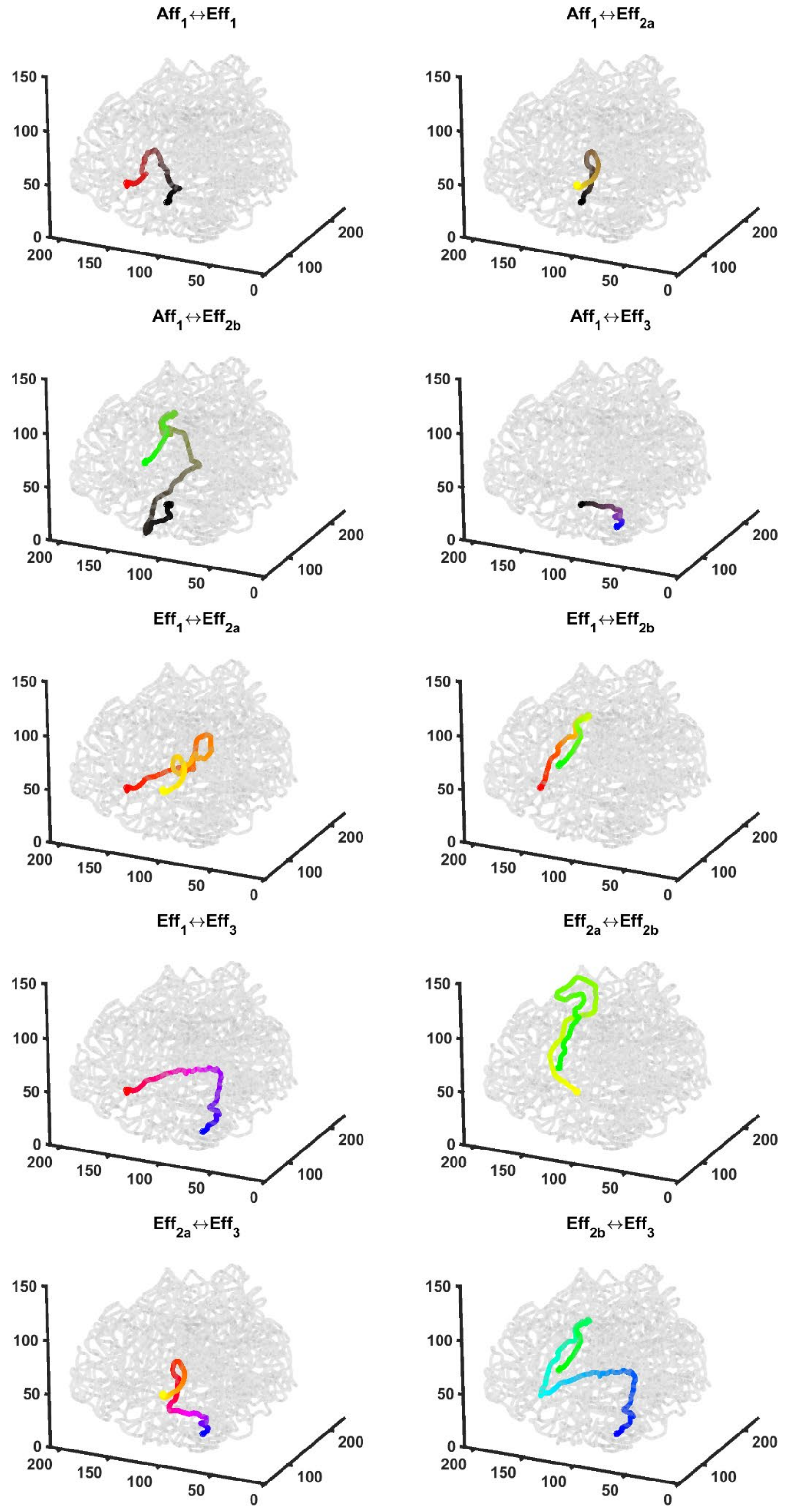


**Supp Fig 4.** Visualization of all pairwise shortest paths between the entry/exit sites. For easy visual reference, the colour coding for $Aff_1$, $Eff_1$, $Eff_{2a}$, $Eff_{2b}$, and $Eff_3$ are black, red, yellow, green and blue, respectively.

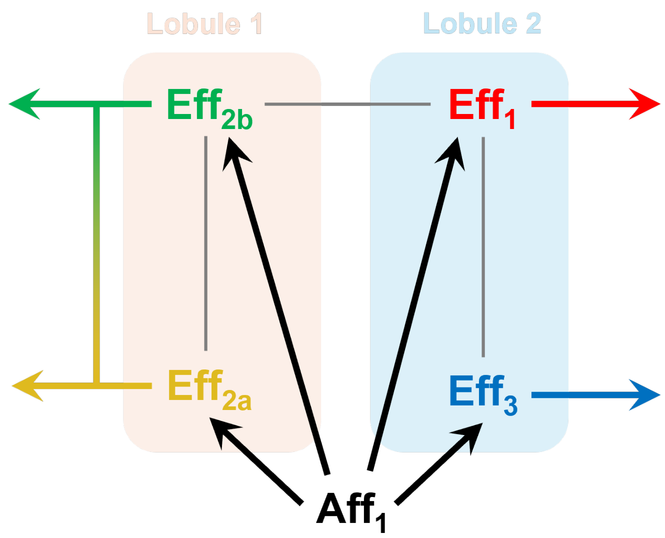


**Supp Fig 5.** Abstracted lobules and domains organization and their connectivities of the glomerulus with multiple efferents, as presented in main **Figures 1** and **2**.

**Supplementary Video**

**Supp Video 1.** 3D image stack rendering of the completely reconstructed human glomerulus as in **Figures 1** and **2**, along with manual tracing of the capillary network and its graph representation.
